# The effects of high-altitude windborne migration on survival, oviposition and blood-feeding of the African malaria mosquito, *Anopheles gambiae* s.l

**DOI:** 10.1101/2020.04.27.065169

**Authors:** ZL Sanogo, AS Yaro, A Dao, M Diallo, O Yossi, D Samaké, BJ Krajacich, R Faiman, T Lehmann

**Affiliations:** Malaria Research and Training Center (MRTC) / Faculty of Medicine, Pharmacy and Odonto-stomatology, Bamako, Mali; Laboratory of Malaria and Vector Research, NIAID, NIH. Rockville, MD, USA

**Keywords:** altitude, blood-feeding, egg-laying, malaria, migration, survival, windborne-dispersal

## Abstract

Recent results of high-altitude windborne mosquito migration raised questions about the viability of these mosquitoes despite ample evidence that many insect species, including other dipterans have been known to migrate regularly over tens or hundreds of kilometers on high-altitude winds and retain their viability. To address these concerns, we subjected wild *An. gambiae* s.l. mosquitoes to a high-altitude survival assay, followed by oviposition (egg laying) and blood feeding assays. Despite carrying out the survival assay under exceptionally harsh conditions that probably provide the lowest survival potential following high altitude flight, a high proportion of the mosquitoes survived for six and even eleven hours assay durations at 120-250m altitudes. Minimal differences in egg laying success were noted between mosquitoes exposed to high altitude survival assay and those kept near the ground. Similarly, minimal differences were found in the female’s ability to take an additional blood meal after oviposition between these groups. We conclude that similar to other high-altitude migrating insects, mosquitoes are able to withstand extended high-altitude flight and subsequently reproduce and transmit pathogens by blood feeding on new hosts.

## Background

The recent report of windborne migrating mosquitoes at high altitude (Huestis *et al.*, 2019) marks a paradigm shift in our understanding of mosquito and pathogen dispersal. Many insect species, ranging in size from large locusts (Orthoptera; Acrididae), hoverflies (Diptera; Syrphidae), blackflies (Diptera; Simulidae), frit flies (Diptera; Chloropidae), wheat midges (Diptera; Cecidomyiidae), and even minute *Culicoides* biting midges (Diptera; Ceratopognidae) and aphids (Hemiptera; Aphididae) have been known to exploit high-altitude winds to migrate over tens or hundreds of kilometers (Johnson *et al.*, 1962; Johnson, 1969; Rainey, 1973; Sellers, 1980; Pedgley *et al.*, 1995; Reynolds *et al.*, 2006; Sanders *et al.*, 2011; Miao *et al.*, 2013; Wotton *et al.*, 2019). However, despite anecdotal observations in support of similar migratory behavior (Glick, 1939; Garrett-Jones, 1962; Reynolds *et al.*, 1996; Johansen *et al.*, 2003), mosquitoes and especially malaria vectors were considered to migrate exclusively in the flight boundary layer, typically well below 10m above ground level (agl) where the mosquito’s own flight is the key factor determining its speed and direction rather than the wind (Snow & Wilkes, 1972; Gillies & Wilkes, 1976; Snow, 1982). Because high-altitude windborne migration in mosquitoes has long been considered accidental and thus of negligible significance (Service, 1997) some vector biologists doubt the viability of the mosquitoes collected in altitude.

In other high-altitude windborne migrant insects, questions about viability post-migration have been settled long ago by studies comparing survival and reproduction in a live collection of insects, including small Diptera (using non-sticky nets, at altitudes similar to our panels) with those captured on the ground or by simulated long flights (Taylor, 1960; Cockbain, 1961; Mcanelly & Rankin, 1986). After finding similar survival and reproductive success, Taylor (1960) concluded that “This seems to establish the viability of high-level migrants beyond reasonable doubt.” The view that insect flight at high altitude is in itself harmful, and insects are subject to physiological stresses not found in flight at low altitude, has become rare (Johnson, 1969), at least among agricultural entomologists. This is partly due to small pest insects evidently migrating over very long distances and infest crops on landing, such as the brown planthopper (*Nilaparvata lugens*) which migrates about 700-1000 km from eastern China to Japan every year (Rosenberg & Magor, 1987). Evidence for the benefit of long-range windborne migration for the insect migrants has also recently came to light based on four fold amplification of the spring migrants when compared with their returning offspring (Chapman *et al.*, 2012).

Considering mosquitoes, specimens caught by aerial netting at altitude in China and India (Ming *et al.*, 1993; Reynolds *et al.*, 1996) were alive and active upon capture. Based on the distinct composition of the mosquito species, sexes, and female gonotrophic states at altitude compared with on the ground, Huestis *et al.* (2019) inferred that mosquitoes, like other insects (Drake & Reynolds, 2012), deliberately ascend into the winds at altitude rather than being inadvertently “forced upwards” by winds. For example, collections 100-290 m agl in Mali were dominated by secondary malaria vectors, e.g., *An. squamosus* and *An. pharoensis*, whereas, on the ground using indoor collections, outdoor clay-pot traps, and larval collections in the vicinity of the same villages, >90% of *Anopheles* captured were *An. gambiae* s.l.. The difference among *An. coluzzii* and *An. arabiensis* that share similar larval, biting, and resting sites (Toure *et al.*, 1996; Lemasson *et al.*, 1997; Lehmann & Diabate, 2008; Dao *et al.*, 2014) and are less affected by sampling bias, better demonstrate species-specific differences in high altitude flight behavior because *An. arabiensis* has not been found at altitude. Additionally, aerial density of mosquitoes was higher when ground-level wind was slower (Huestis *et al.*, 2019; Florio *et al.*, 2020: PREPRINT).

In addition to the exertion of sustained flight, presumably over several hours (Kaufmann & Briegel, 2004; Huestis *et al.*, 2019; Faiman *et al.*, 2020: PREPRINT), nightly high altitude flight exposes mosquitoes to a combination of different temperatures, humidity (RH) and wind speeds than those conditions on the ground. Given the low sampling efficiency of mosquitoes in high altitude, evaluating the effects of these factors on their viability is not straightforward. In a preliminary analysis described in Huestis *et al.* (2019), survival of *Anopheles gambiae* s.l. collected indoors and placed individually, in modified 50 ml tubes (both ends covered with netting, Fig S1) that were raised using the helium balloon to 120-190 m agl and subjected to wind passing through the tubes for 13 hours was not statistically different from that of mosquitoes kept near the ground (altitude: 58% N=26 vs. ground: 71%, N=17; P>0.38, χ^2^_1_=0.75, Fig. 1a). Given that the mosquitoes at altitude were unable to “ride the wind” but were tumbling against and abraded by the hard stretched net all night long, this assay provides the lowest survival limit of mosquitoes at altitude. Nonetheless, without a better alternative, here we utilized this conservative assay to measure the effect of altitude, duration of “flight”, and wind speed on the mosquito’s survival. Additionally, we evaluate her post-flight capacity to lay eggs, and her ability to take another blood meal. Our new results, based on a larger sample size, demonstrate that mosquito migrants at high altitude can indeed survive, lay eggs, and thereafter take a new blood meal, thus enabling a new transmission encounter with the host after their migration.

**Fig. 1.**
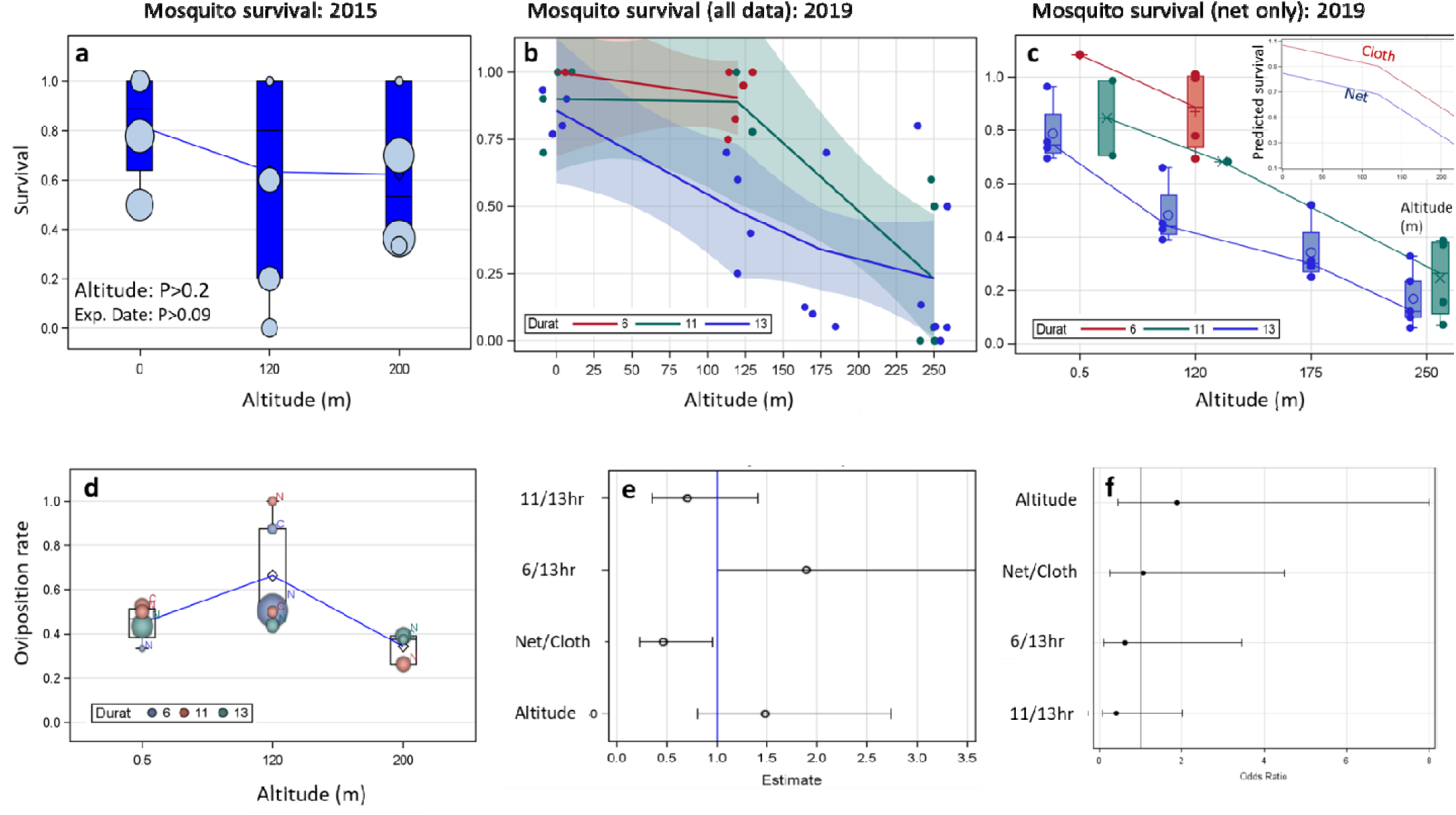
The effect of altitude and assay duration on survival of A. gambiae s.l. mosquitoes. (a) The survival rates in July and October 2015 for 13 hours assay duration described in Huestis *et al.* (2019) and Introduction (above); dot size signifies sample size per experimental date (lines connect the mean values). (b) The survival rate in late October-November 2019 over different assay durations (pooling net and cloth covers of the tubes). (c) The survival rate (2019) over different assay times based on least square means (net only). Inset: The difference in survival between tubes covered with net (blue) or cloth (red) for the means of the 11-13 hours exposure assays. d) Oviposition rate (2019) among survivors from different altitudes, assay durations and net covers (N vs. C). e) Odds ratios estimates (dot) and 95% CI of the probabilities to lay eggs between treatments denoted on the Y axis based. Note: if the 95% CI intersects 1, the effect is not statistically significant. f) Odds ratios estimates (dot) and 95% CI of the probability to blood feeding (after oviposition, see e, above).

## Methods

### Study location and mosquitoes

This study was performed from October to November 2019 in Thierola, Mali (13°39⍰30.96⍰⍰ N, 7°12⍰52.92⍰⍰ W), a Sahelian village described previously (Lehmann *et al.*, 2010; Dao *et al.*, 2014). The area’s single wet season occurs between June and October (∼550 mm). Rainfall is negligible (<50 mm) from November until May. By December, water is available only in deep wells.

Wild *An. gambiae* s.l. females were collected in human dwellings between 08:00 to 10:00 in Thierola and the neighboring villages (<7 km away) using mouth aspirators. Female mosquitoes were provided with 10% sucrose solution in cages covered with wet towels, which were kept in a typical village house used as a field insectary (without climate control). Blood-fed and semi-gravid female mosquitoes were housed in the field insectary until they reached the gravid state (up to 2 days) and could be subjected to the high-altitude survival assay. Females had access to 10% sugar solution until 2 hours before the survival assay.

### The high-altitude survival assay

Fully gravid females were randomly assigned to different altitude exposure treatments varying between a) 1 to 290 m agl, b) assay duration of 6, 11, or 13 hours, and c) high vs. low air flow. Each female was individually placed in 5 cm long and 3 cm diameter tubes made by cutting 50 ml Falcon tubes (Fig. S1). To control the air flow through the tubes, the openings were covered with net (hole diameter= 1.5mm) or cloth (hole diameter= 0.2mm, Fig. S1). Groups of mosquitoes were launched after sunset and retrieved around sunrise, except the six hour duration group, which were either launched and retrieved between 18:00 and midnight or between midnight and 06:00 as previously described (Huestis *et al.*, 2019). Five to ten tubes containing mosquitoes were mounted on the rope using adhesive tape (Fig. S1) in set altitudes 1, 120, 180 (160 and 190 pooled) and 250 (220-280 pooled) m from the ground. Mosquitoes mounted 1 m from the ground and those kept in insectary were used as controls. Upon retrieval, typically around 07:00, mosquitoes were examined for mobility and recorded as live (mobile) or dead (immobile) within one hour after retrieval. Live mosquitoes were further subjected to oviposition assay.

### Oviposition assay

Surviving mosquitoes were individually transferred into 50 ml tube with 5 ml water for oviposition on the afternoon of the same day they completed the survival assay. Every morning, during four consecutive days, each tube was inspected for eggs. The number of eggs laid was estimated and their hatching was noted in the following days. Females that died during the oviposition assay were scored to produce zero eggs. Females that did not lay eggs by the end of the oviposition assay were killed and immediately dissected and their spermatheca examined to determine their insemination status. Their ovaries were also examined to determine if they were gravid and the number of developed eggs in their abdomen were counted. Due to logistical constraints, not all females that did not lay eggs were dissected.

### Blood feeding assay

Females which laid eggs were subjected to a blood feeding assay the following night. They were provided with water only (no sugar solution) until 22:00, when they were placed, in a pint size cage, against a chicken’s breast (under the wing) of an immobilized chicken for 20 min in accord with animal care guidelines (F20-00465 MRTC). Immediately afterwards, females were scored as fully fed, partly fed, or unfed.

At the end of the blood feeding assay or after female mosquitoes died naturally (or accidentally), they were preserved in 80% ethanol. The sibling species of the Anopheles gambiae complex were identified as previously described (Fanello *et al.*, 2002).

### Statistical analysis

Mosquito survival, oviposition, and subsequent blood feeding are dichotomous variables. Their corresponding fractions in each category was computed and plotted. To increase group size and the power of the statistical analyses, adjacent altitudinal panels were pooled together, e.g., 160m and 190m were pooled together in a class of 175m and 220-280m were similarly pooled into 250m class. Likewise, we pooled groups of mosquitoes that were exposed to altitude between 18:00 and midnight with those that were exposed from midnight to 06:00 in the 6-hour duration group. Contingency tables and log-likelihood tests were used to examine the relationship of each treatment separately on the dependent variables (survival, oviposition, and blood feeding), including stratification across an additional variable using Cochran-Mantel-Haenszel test (SAS Inc., 2012). Multivariate analysis of the survival rate of mosquitoes was carried out using Proc Mixed (SAS Inc., 2012) on the fraction of surviving mosquitos per treatment (combination of altitude, duration, cover type and date). Date was introduced in the model as a random variable because it captures variation in temperature, wind speed, and RH (below). To evaluate the variation among species, the analysis was repeated with and without the species effect. Finally, the nightly weather parameters were introduced into the model. Multivariate analyses of oviposition (egg laying) and blood feeding were carried out using logistic regression carried out by Proc Logistic (SAS Inc., 2012). Weather data including hourly temperature, RH, wind speed, and direction at 2m and 180m agl were extracted from atmospheric reanalyses of the global climate ERA5 (Copernicus Climate Change Service, 2018) as previously described (Huestis *et al.*, 2019). Nightly means of each parameter from 18:00 to 07:00 (Fig. S2) at corresponding experimental nights were used as predictors of mosquito survival (Table 1).

**Table 1.** Distribution of An. gambiae s.l. across treatments and their survival rate

| Altitude | Duration (h) | Cover | N | Survival (%) |
| --- | --- | --- | --- | --- |
| Ground | 6 | Net | 7 | 100 |
| Ground | 6 | Cloth | ND | ND |
| Ground | 11 | Net | 20 | 85 |
| Ground | 11 | Cloth | 20 | 95 |
| Ground | 13 | Net | 58 | 85 |
| Ground | 13 | Cloth | ND | ND |
| 120m | 6 | Net | 127 | 91 |
| 120m | 6 | Cloth | 10 | 100 |
| 120m | 11 | Net | 9 | 78 |
| 120m | 11 | Cloth | 10 | 100 |
| 120m | 13 | Net | 33 | 49 |
| 120m | 13 | Cloth | ND | ND |
| 175m | 13 | Net | 65 | 19 |
| 250m | 11 | Net | 66 | 35 |
| 250m | 11 | Cloth | 10 | 0 |
| 250m | 13 | Net | 74 | 12 |
| 250m | 13 | Cloth | 10 | 80 |

## Results & Discussion

### Survival after high altitude exposure assay

Over nine nights, a total of 519 wild *An. gambiae* s.l. females were subjected to the survival assay (Fig. S1, Methods) and maintained for 6 to 13 hours in altitudes ranging from 1 to 280 m agl (Table 1). Because wind speed increases with altitude and the mosquitoes remain confined in their tubes against the wind, rather than fly almost stationary in relation to the parcel of air they would be carried by, we predicted that most stressful conditions occur in the longest assay at the highest altitude in tubes covered by net vs. cloth.

Overall, the difference in survival due to altitude varied little between ground (91%, n=105) and 120m (84%, n=188), but was large at higher altitudes (160-280m: 25%, n=225, Table 1). As expected, survival fell with exposure time: 92% (n=144), 56% (n=135) and 39% (n=240) for 6, 11, and 13 hours, respectively. Additionally, survival at altitude increased if the openings of the tubes were covered by a cloth of higher wind resistance (0.2mm hole sizes: 78%, n=60) compared with tubes covered by net of lower wind resistance (1.5mm hole size: 56%, n=411). However, because the experiments were not balanced, the similar survival rate between the ground and 120 m altitude compared with the lower survival at 160--280m, probably was affected by the inclusion of short-duration exposure (6 hours and to a lesser extent 11 hours) at 120m agl (Fig 1). To parse these effects, we analyzed them simultaneously using ANCOVA with random variable (date) and fixed effects of duration, altitude, and net type. As expected, the results revealed that altitude, assay duration, and wind-resistance of the tube cover had significant effect on mosquito survival (P<0.027, Table 2), whereas the variance among dates was non-significant (P<0.067, Table 2). On average, 100m increment of altitude was associated with 25% reduction in survival and an additional hour of the assay reduces survival by 6% (Table 1). Using higher wind resistant cover (cloth) over the tube’s opening instead of lower wind-resistant netting increased survival by 22% (Table 2). The species composition at the time of the assay was dominated by *A. coluzzii* (76%), followed by *A. arabiensis* (14%) and *A. gambiae s.s.* (10%, N=344 mosquitoes). The variation among the species in survival was not significant (P>0.35, Table 2) and the other effects remained unchanged, indicating that the three Sahelian species responded similarly to the assay.

**Table 2.**
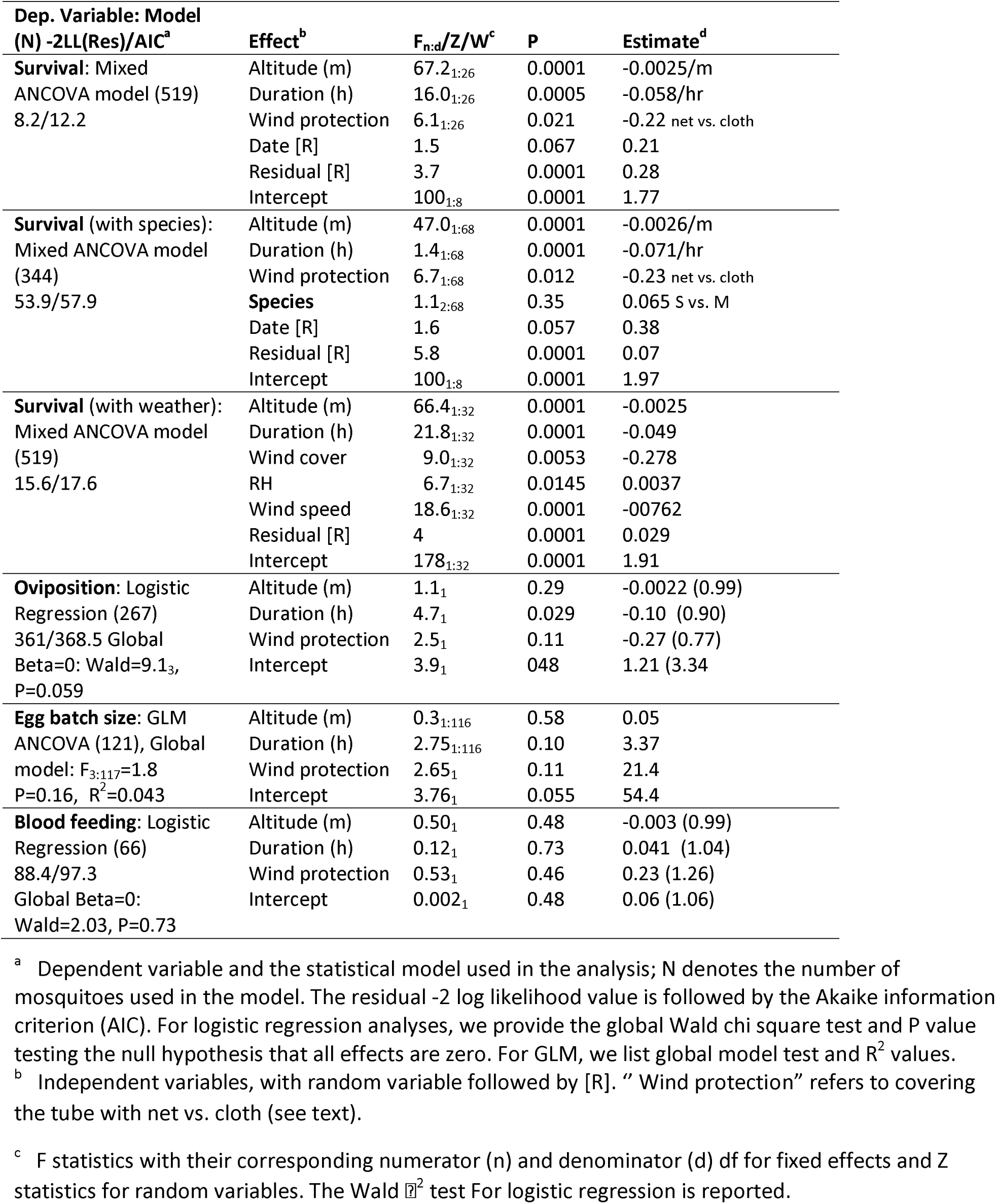
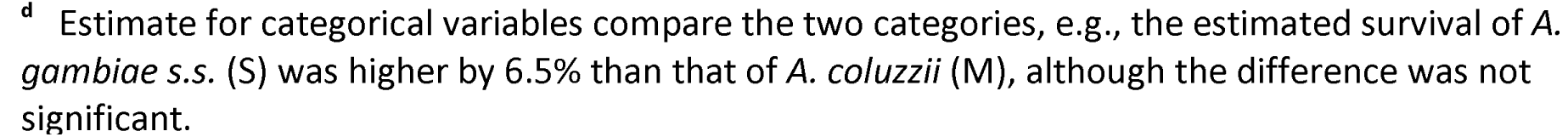
Summary of the results of the statistical models used to analyze survival, oviposition success, and blood feeding success.

| Dep. Variable: Model<br>(N) -2LL(Res)/AIC<sup>a</sup> | Effect<sup>b</sup> | F<sub>n,d</sub>/Z/W<sup>c</sup> | P | Estimate<sup>d</sup> |
| --- | --- | --- | --- | --- |
| **Survival:** Mixed ANCOVA model (519)<br>8.2/12.2 | Altitude (m) | 67.2<sub>1:26</sub> | 0.0001 | -0.0025/m |
|  | Duration (h) | 16.0<sub>1:26</sub> | 0.0005 | -0.058/hr |
|  | Wind protection | 6.1<sub>1:26</sub> | 0.021 | -0.22 net vs. cloth |
|  | Date [R] | 1.5 | 0.067 | 0.21 |
|  | Residual [R] | 3.7 | 0.0001 | 0.28 |
|  | Intercept | 100<sub>1:8</sub> | 0.0001 | 1.77 |
| **Survival** (with species): Mixed ANCOVA model (344)<br>53.9/57.9 | Altitude (m) | 47.0<sub>1:68</sub> | 0.0001 | -0.0026/m |
|  | Duration (h) | 1.4<sub>1:68</sub> | 0.0001 | -0.071/hr |
|  | Wind protection | 6.7<sub>1:68</sub> | 0.012 | -0.23 net vs. cloth |
|  | **Species** | 1.1<sub>2:68</sub> | 0.35 | 0.065 S vs. M |
|  | Date [R] | 1.6 | 0.057 | 0.38 |
|  | Residual [R] | 5.8 | 0.0001 | 0.07 |
|  | Intercept | 100<sub>1:8</sub> | 0.0001 | 1.97 |
| **Survival** (with weather): Mixed ANCOVA model (519)<br>15.6/17.6 | Altitude (m) | 66.4<sub>1:32</sub> | 0.0001 | -0.0025 |
|  | Duration (h) | 21.8<sub>1:32</sub> | 0.0001 | -0.049 |
|  | Wind cover | 9.0<sub>1:32</sub> | 0.0053 | -0.278 |
|  | RH | 6.7<sub>1:32</sub> | 0.0145 | 0.0037 |
|  | Wind speed | 18.6<sub>1:32</sub> | 0.0001 | -0.0762 |
|  | Residual [R] | 4 | 0.0001 | 0.029 |
|  | Intercept | 178<sub>1:32</sub> | 0.0001 | 1.91 |
| **Oviposition:** Logistic Regression (267)<br>361/368.5 Global<br>Beta=0: Wald=9.1<sub>3</sub>,<br>P=0.059 | Altitude (m) | 1.1<sub>1</sub> | 0.29 | -0.0022 (0.99) |
|  | Duration (h) | 4.7<sub>1</sub> | 0.029 | -0.10 (0.90) |
|  | Wind protection | 2.5<sub>1</sub> | 0.11 | -0.27 (0.77) |
|  | Intercept | 3.9<sub>1</sub> | 0.048 | 1.21 (3.34) |
| **Egg batch size:** GLM ANCOVA (121), Global model: F<sub>3,117</sub>=1.8<br>P=0.16, R<sup>2</sup>=0.043 | Altitude (m) | 0.3<sub>1:116</sub> | 0.58 | 0.05 |
|  | Duration (h) | 2.75<sub>1:116</sub> | 0.10 | 3.37 |
|  | Wind protection | 2.65<sub>1</sub> | 0.11 | 21.4 |
|  | Intercept | 3.76<sub>1</sub> | 0.055 | 54.4 |
| **Blood feeding:** Logistic Regression (66)<br>88.4/97.3<br>Global Beta=0:<br>Wald=2.03, P=0.73 | Altitude (m) | 0.50<sub>1</sub> | 0.48 | -0.003 (0.99) |
|  | Duration (h) | 0.12<sub>1</sub> | 0.73 | 0.041 (1.04) |
|  | Wind protection | 0.53<sub>1</sub> | 0.46 | 0.23 (1.26) |
|  | Intercept | 0.002<sub>1</sub> | 0.48 | 0.06 (1.06) |

<sup>a</sup> Dependent variable and the statistical model used in the analysis; N denotes the number of mosquitoes used in the model. The residual -2 log likelihood value is followed by the Akaike information criterion (AIC). For logistic regression analyses, we provide the global Wald chi square test and P value testing the null hypothesis that all effects are zero. For GLM, we list global model test and R<sup>2</sup> values.
<sup>b</sup> Independent variables, with random variable followed by [R]. "Wind protection" refers to covering the tube with net vs. cloth (see text).
<sup>c</sup> F statistics with their corresponding numerator (n) and denominator (d) df for fixed effects and Z statistics for random variables. The Wald $\chi^2$ test For logistic regression is reported.
<sup>d</sup> Estimate for categorical variables compare the two categories, e.g., the estimated survival of *A. gambiae* s.s. (S) was higher by 6.5% than that of *A. coluzzii* (M), although the difference was not significant.

Under this conservative survival assay, females of *A. coluzzii, A. gambiae* and *A. arabiensis* can survive >13 hours at high altitude. Highest survival (>90%) was shown when exposure was 6 hours or up to 11 hours at 120m when the wind force was attenuated by a cloth (pores of 0.2mm diameter) instead of a net (pores of 1.5mm diameter). Mean nightly windspeed at 150m agl (5--7m/s) was >5 fold greater than at 2m agl (Fig. S2), and >7 fold at 250m agl (7--9m/s, not shown), explaining the harsh conditions mosquitoes experience in tubes covered by nets. The small pore size of the cloth allows rapid equilibration of temperature and RH with the surroundings, so the protective effect of the cloth operates solely by wind attenuation. This is also corroborated by the negative effect of altitude on survival because wind speed increases with altitude. The survival assay is extremely harsh because the mosquito is pummeled by strong wind against the rough surface of the stretched net, probably resulting in desiccation and physical damage that increase mortality. This effect does not occur in natural high-altitude flight when the mosquito is more or less stationary with respect to the air parcel it is carried in. Additionally, the 2019 experiments were performed in the transition between the wet (October) and the dry season (November) when nightly relative humidity drops from 80% to 30% (Fig S2) and mean nightly windspeed at altitude increased during November to 9-10 m/s compared with 5-6 m/s in August-September (Fig. S2), during peak migration (Huestis *et al.* 2019). *Importantly, even under these exaggerated, taxing conditions, 70% and 30% of the mosquitoes survived for 11 hours at 120 or 250 m agl, respectively as opposed to 90% at ground level*.

### Oviposition of mosquitoes that survived high-altitude exposure assay

To assess if gravid *A. gambiae* s.l. mosquitoes that withstood exposure to high altitude (above) are capable of laying eggs, they were transferred to individual 50 ml tubes and provided with water for oviposition. The water was examined daily for eggs during four days. Mosquitoes that did not lay eggs were dissected to determine insemination status. Overall, 46% of the 267 gravid females subjected to the assay laid eggs. However, 12% (n=10) of 83 females that did not lay eggs and were dissected had no sperm and therefore, could not lay eggs, thus indicating that overall oviposition rate among inseminated females was near 60% (the average egg batch size was 108 (n=121, 95% CL=97-119). During October November oviposition rate and egg batch size of A. coluzzii are reduced (Yaro *et al.*, 2012), and the observed values are similar to those observed previously during this time.

The effects of altitude, assay duration, and tube cover material on the likelihood of laying eggs were weak (Table 2, Fig. 1d and Fig. 1e). The effect of altitude was not significant (P>0.2, Table 2, Fig. 1e). The effect of the assay duration was significant (P<0.03, Table 2), but the 95% confidence limits of the highest odds ratio (6 vs. 13 hours) included 1 (Fig. 1e), questioning the significance of the effect. The effect of tube cover was statistically significant (P<0.039, Table 2) and amounted for 32% higher egg laying probability compared with those housed in a net covered tube (Table 2). Considering egg batch size of females that laid eggs (excluding zeros, N=121), the effects of altitude, assay duration, and tube cover material were all not significant (Table 2). For example the mean egg batch size (and 95%CI) of females kept on the ground, at 120m, and 200m agl were 110.5 (92.9-128.0, N=39), 108.9 (93.7-122.5, N=66), and 101.7 (62.4-141.0, N=16), respectively and the largest egg batch size (340 eggs) was laid by a female kept at 200 m agl.

### Blood feeding of mosquitoes that survived high-altitude exposure assay

To assess if gravid A. gambiae s.l. mosquitoes that withstood exposure to high altitude (above) are capable of taking a blood meal after laying eggs, females were subjected to a blood feeding assay (see Methods). Overall, 56% of the 66 females subjected to the assay took a blood meal. Differences between treatments were minimal and statistically not significant (Table 2. Fig. 1f). The rates of blood feeding on the ground vs. at altitude were 65% and 50%, respectively; at assay times of 6, 11, and 13 hours were 52%, 50%, and 73%, respectively; and under net vs. cloth were 58% and 50% respectively.

## Conclusions

These experiments extend previous results obtained in 2015 (Huestis *et al.* 2019). In addition to larger sample size for the survival analysis, surviving mosquitoes were subjected to an oviposition assay followed by a blood feeding assay to evaluate the capacity of anopheline mosquitoes to survive the exposure to high altitude, and subsequently to lay eggs and take, at least, one additional blood meal. Despite carrying out these experiments during the transition from the wet to the dry season (October--November), after peak migration (Huestis *et al.*, 2019), when RH decreases and wind speed increases – conditions that reduce mosquito survival (Clements, 1992; Huestis & Lehmann, 2014; Arcaz *et al.*, 2016) and despite using an exceptionally harsh survival assay that arguably provides the lowest limit of survival following high altitude flight, high proportion of the mosquitoes survived for 11 hours assay duration. Furthermore, minimal differences in egg laying and in their ability to take another blood meal were found between mosquitoes exposed to high altitudes overnight and those near the ground. We conclude that similar to all other insect species that have been evaluated (Taylor, 1960; Cockbain, 1961; Mcanelly & Rankin, 1986) mosquitoes are able to withstand high altitude flight and subsequently reproduce and transmit pathogens by blood feeding on new hosts.

## Acknowledgements

We are grateful to the residents of Thierola for their permission to work near their homes and for their wonderful assistance and hospitality. We thank Drs. Sekou F Traore and Thomas Wellems, and Ms. Margie Sullivan, and Mr. Samuel Moretz (National Institutes of Health, USA) for logistical support. Dr. Malla Rao (NIH) has highlighted the importance of the question addressed by this study to the vector biology community in a discussion with Dr. Yan Guiyun (University of California, Irvine). We thank Drs. Don Reynolds (University of Greenwich, UK) and Jason Chapman (University of Exeter, UK and Nanjing Agricultural University, China) for reading earlier versions of this manuscript and providing us with helpful suggestions.

## Funding

This study was supported by the Division of Intramural Research, National Institute of Allergy and Infectious Diseases, National Institutes of Health.

## Authors Contributions

This project was designed by TL, AD, ASY, and RF. Field methods and operations, data management, and specimens processing was performed by ZLS, ASY, MD, DS, and OY. Laboratory processing and identification of morphotypes was done by ZLS, MD, DS, OY, ASY, and AD. Data analysis were carried out by TL with inputs from all authors, especially BJK, RF, ZLS and ASY. BJK has written scripts to obtain ERA5 data and provided these data. The manuscript was drafted by TL and ZLS and revised by all authors. Throughout the project, all authors have contributed key ingredients and ideas that have shaped the work and the final paper.

## Competing Interests

All authors declare no competing financial interests.

## Data availability

The dataset and SAS code used to analyze the data are available upon request from the corresponding author.

## Notes

### Competing Interest Statement

The authors have declared no competing interest.

