## Supplemental Figures 1 and 2 for "The effects of high-altitude windborne migration on survival, oviposition and blood-feeding of the African malaria mosquito, *Anopheles gambiae* s.l"

12735 Twinbrook Parkway, Room 3W-13-F

Rockville MD 20852	Cell :    240 408 9820

**Fig. S1**. Tubes used to house mosquitoes during the survival at altitude assay. A standard 50 ml tube was cut in half to form two experimental tubes (after removal of the tapering end, Methods). a) design highlighting net vs. cloth covers of the openings. b) Tubes mounted on the rope tethered between the ground and the helium balloon. c) Tubes in a tray with mosquitoes inside (arrows).

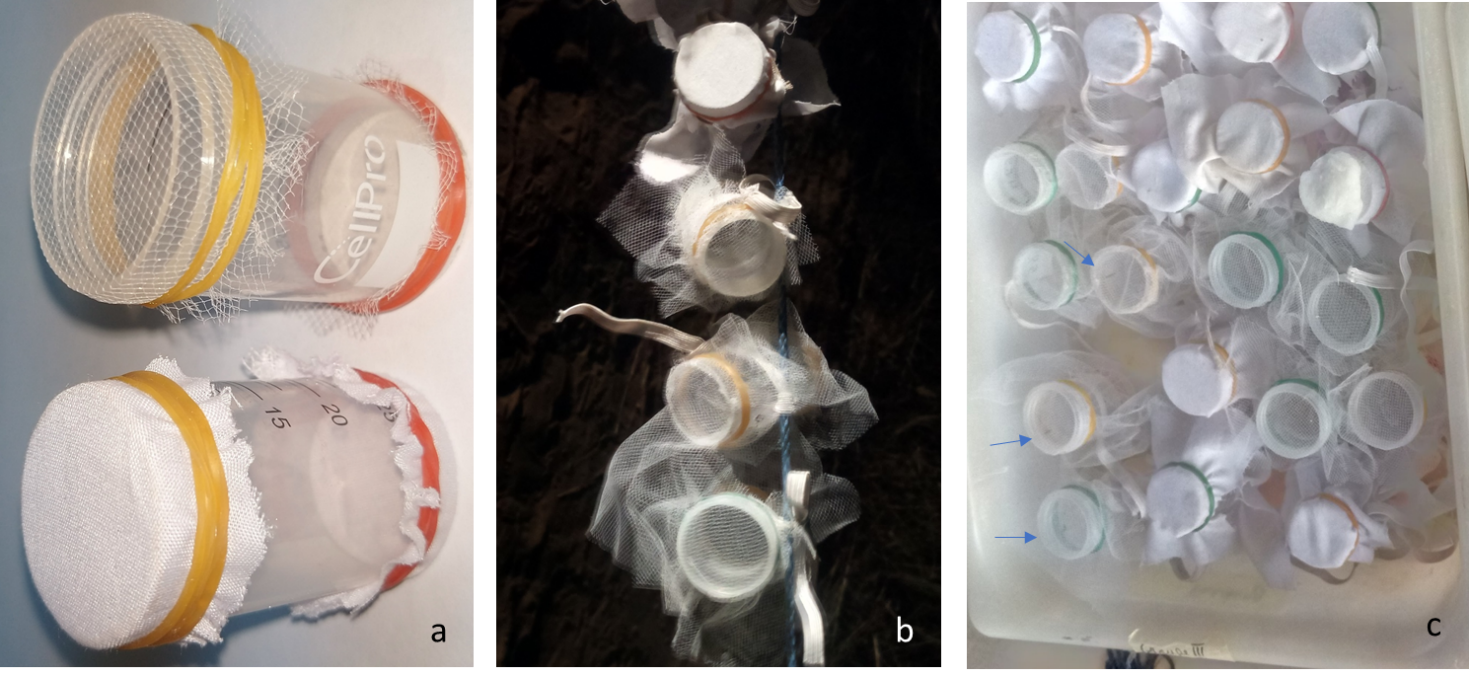

**Fig. S2**. Seasonal change in nightly (17:30 through 07:30) RH (a and d), wind speed (b and e), antemperature (c and f) in Thierola during the experiment (October-November 2019) at 145 m (source: ERA5, a, b, and c) and 2 m agl (d, e, and f; source: Rainwise Inc. weather station). Dots represent hourly records (145m agl) and 30 min (2 m agl). Lines represent trends based on local regression (loess). Red ‘v’ signify nights when the survival assay were carried out.

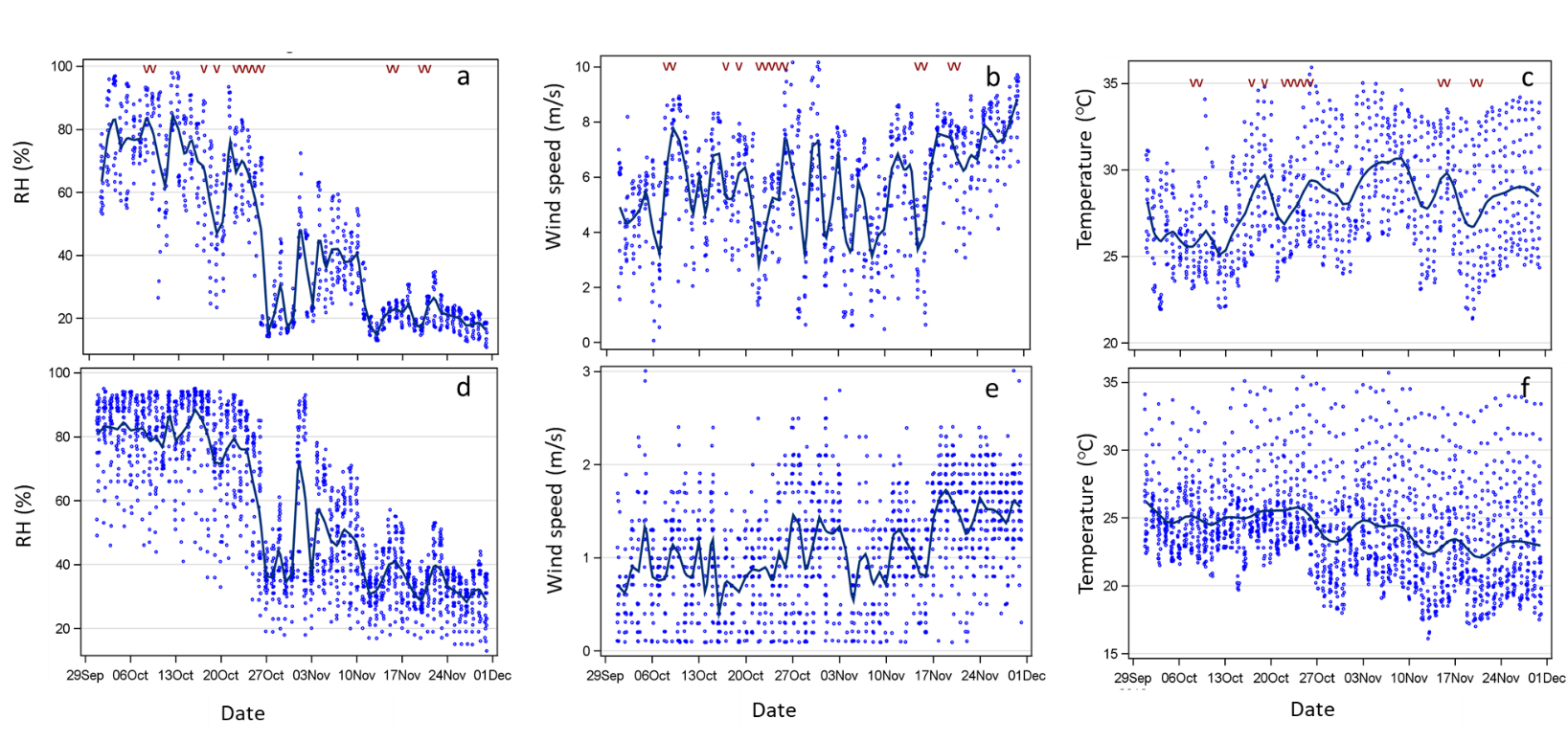
